# Post-encoding administration of oxytocin selectively enhances memory consolidation of male faces in females

**DOI:** 10.1101/2024.06.18.599465

**Authors:** Jiashen Li, Zhengyue Chen, Wei Liu

**Author notes:** Correspondence: Dr. Wei Liu, School of Psychology, Central China Normal University, Address: the 8^th^ floor, Nanhu Complex Building, No 152 Luoyu Road, Wuhan. Postal Code: 430079.

## Abstract

Oxytocin plays a critical role in modulating social cognition and enhancing human memory for faces. However, it remains unclear which stage of memory oxytocin affects to enhance face memory. Our study explored oxytocin’s potential to selectively enhance the consolidation of social memories, specifically human faces, and whether this effect varies between genders. In two preregistered, randomized, double-blind, placebo-controlled trials with heterosexual participants (total N=294, comprising 149 males and 145 females), we explored how oxytocin affects memory consolidation. We administered oxytocin immediately after encoding (i.e., Study1) and 30 minutes before retrieval in a parallel study (i.e., Study2). This design allowed us to confirm that oxytocin’s effects were indeed due to consolidation rather than retrieval. We found that administering oxytocin post-encoding, but not before-retrieval, significantly improved female participants’ ability to recognize male faces 24 hours later, with no similar enhancement observed in males recognizing opposite-gender faces. Together with our analyses of social placebo effects and approachability rating during encoding, we concluded that oxytocin enhanced consolidation of long-term social memories in humans. Our results not only advance the understanding of the neurobiological mechanisms underlying social memory consolidation but also highlight oxytocin as a pharmacological tool for selectively enhancing human memory consolidation.

**Highlights:**

1. Oxytocin selectively enhances memory consolidation of human faces, with gender-specific effects.
2. In females, oxytocin after encoding improves recognition of male faces after 24 hours.
3. Oxytocin-induced enhancement of social memory is due to enhanced consolidation, not retrieval or encoding.
4. Oxytocin shows potential for selectively modulating memory consolidation in humans.

## Introduction

From rodents to non-human primates and humans, social cognition plays a critical role in building and maintaining social relationships. Studies from the last two decades have demonstrated the modulatory role of neuropeptide oxytocin in social cognition from basic (e.g., modulate social behaviors such as competition and cooperation and their underlying neural networks^1–3^) to clinical science (e.g., potential treatment of autism^4^, psychopathy^5^, and social anxiety disorder^6^). This field of intranasal oxytocin research also starting to shed light on the debatable question that emerged in the cross between social cognition and memory: whether social memory (e.g., faces) and other forms of memory (e.g., houses) have separate neural and biological basis? In rodents, oxytocin knock-out nice showed impaired social memory, but intact nonsocial memory^7^. Critically, the impairment in social memory could be fully restored by a single injection of oxytocin before the social encounter^8^. In humans, Rimmele et al. showed that oxytocin, administrated before memory encoding, specifically improves recognition memory for faces (but not for non-social stimuli), which was tested 24 hours later^9^. However, the Rimmele’s study was not able to dissociate the effect of oxytocin on memory encoding and consolidation underlying the reported memory enhancement. More specifically, in Rimmele’s study, because the oxytocin was administrated before learning, and memory was tested twenty-four hours later, there is a possibility that the oxytocin simultaneously enhanced memory encoding and consolidation of social memories, leading to improved long-term memories.

To isolate the effects of oxytocin on memory consolidation, we used a post-study design (i.e., oxytocin administrated after memory encoding) based on memory consolidation studies in both animals and humans^10,11^. This is the standard paradigm to evaluate and isolate the effects of certain experimental manipulations (e.g., physical exercise^12^, caffeine intake^13^, brain stimulation^14^, and emotional arousal^15,16^) on memory consolidation. We hypothesized and examined the premise that oxytocin, a pivotal neuropeptide in social cognition, might selectively influence the consolidation of social memories. Previous studies on human memory consolidation suggest that while it can be modulated, this modulation is largely non-selective. Targeted memory reactivation (TMR)^17,18^ is an exception, as it can selectively reactivate memories during sleep—a crucial phase for consolidation. In summary, previous techniques for manipulating memory consolidation in humans have either lacked specificity or depended on particular stages of sleep. This raises the question: Is it possible to develop novel methods for selectively modulating memory consolidation in humans? Furthermore, based on the interaction between oxytocin and social memory, can oxytocin be used as a pharmacological tool to selectively enhance social memory consolidation in humans?

## Results

We conducted two preregistered, randomized, double-blind, placebo-controlled trials with heterosexual participants (N=294) to determine whether oxytocin can selectively affect social memory consolidation in humans. We implemented the recommendations for enhancing future intranasal oxytocin research set forth by Quintana and colleagues^1^: (1) We preregistered our data acquisition and analysis protocols prior to collecting data, minimizing p-hacking and the chance of false-positive results; (2) We intentionally recruited male (n=149) and female (n=145) participants to explore if the selective enhancement of social memory observed in Rimmele’s study^9^ could be extended to heterosexual women, especially given that oxytocin’s cognitive effects can be sex-specific^19–21^. This approach addresses a gap in the literature as most prior studies using oxytocin, including Rimmele’s, predominantly featured male subjects; (3) We made our experimental materials, code, open data, and analysis scripts publicly accessible to facilitate direct replications and independent verification of our statistical findings. (Note: the links to preregistration protocals and shared materials could be found in the STAR Methods section)

Our experimental protocol spanned two sessions conducted over consecutive days, separated by a 24 hours interval. On the first day, participants incidentally encoded images of social (i.e., human faces) and non-social (i.e., houses) stimuli, followed by administration of either oxytocin or a placebo. The timing of administration was immediate—post-encoding in Study1 and pre-retrieval in Study2. After a day, we unexpectedly evaluated participants’ memory retention using the remember/know (RK) paradigm^22^. Memory assessment involved computing the overall recognition score, derived from the overall hit rate minus the false alarm rate, to quantify memory accuracy for various stimuli, including faces and houses (**Figure 1**).

**Figure 1:**
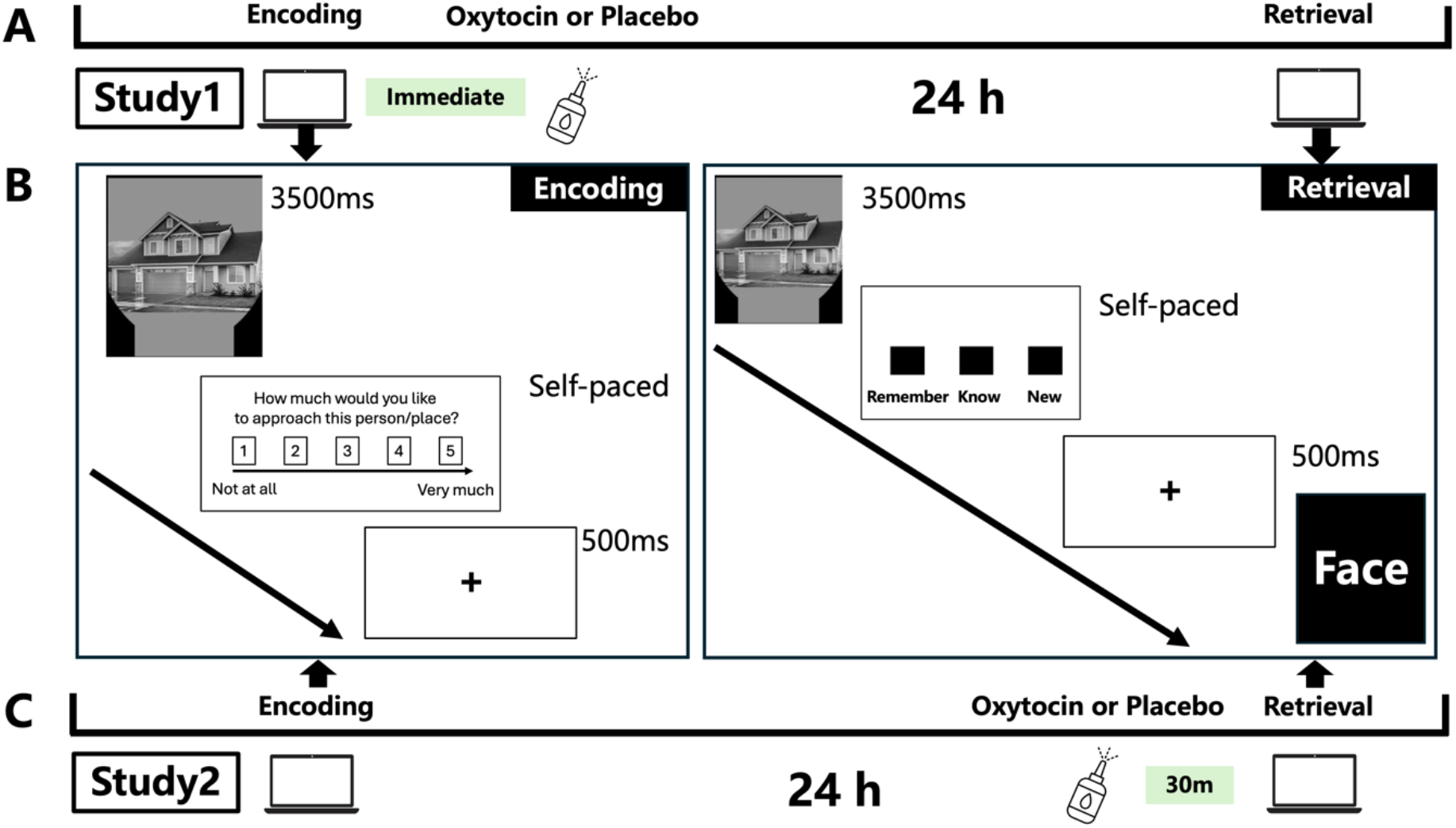
Study Design and Experimental Procedure. **(A) Outline of Study 1:** Participants who passed the screening were immediately administered a 24 IU dose of oxytocin or a placebo following the encoding task. They returned 24 hours later for the retrieval phase, which included a Recognition Test. **(B) Trial Structure for Encoding and Retrieval Tasks:** During the encoding phase, participants engaged in an incidental approachability rating task. Each trial started with the presentation of a picture (i.e., face or hourse) for a specified duration (i.e., 3500 ms), followed by a self-paced rating on an approachability scale (ranging from 1, not at all, to 5, very approachable). An interstimulus interval (ISI) of 500 ms separated each trial. In the retrieval phase, recognition was assessed using a Remember/Know (RK) paradigm with targets and similar lures. Each picture was displayed for 3.5 seconds, with a 0.5-second ISI. Participants indicated whether they remembered (R), knew (K) the stimulus, or considered the picture new by pressing one of three response keys. **(C) Outline of Study 2:** Study 2 employed the same encoding and retrieval tasks as Study 1, with the only modification being the timing of oxytocin or placebo administration, which was adjusted to 30 minutes before retrieval, approximately 23.5 hours after encoding. Note: *We used real faces from the Chinese Academy of Sciences’ Sinicized Face Emotion Image System. In this figure, we used the word ’face’ instead of actual face examples to avoid displaying identifiable information from human volunteers*.

### Gender-specific enhancement of social memory consolidations in humans

During data collection for Study1, we strictly adhered to our pre-registered protocol, with two deviations: (1) an extended data acquisition period beyond the preregistered duration due to slow progress; (2) a sample size slightly differing from the target of 30 per group, ranging from 29 to 33 participants (see **Table 1** for exact sample sizes and memory performance per group). The sample size deviation occurred because we could not accurately predict participant exclusions based on the pre-registered data analysis protocol, which excluded participants with low memory performance (i.e., overall recognition score lower than 0). We also conducted exploratory analyses beyond preregistered protocal (e.g., included all tested pariticpants in data analyses regardless of memory performance), which are reported separately in this manuscript.

**Table 1.**
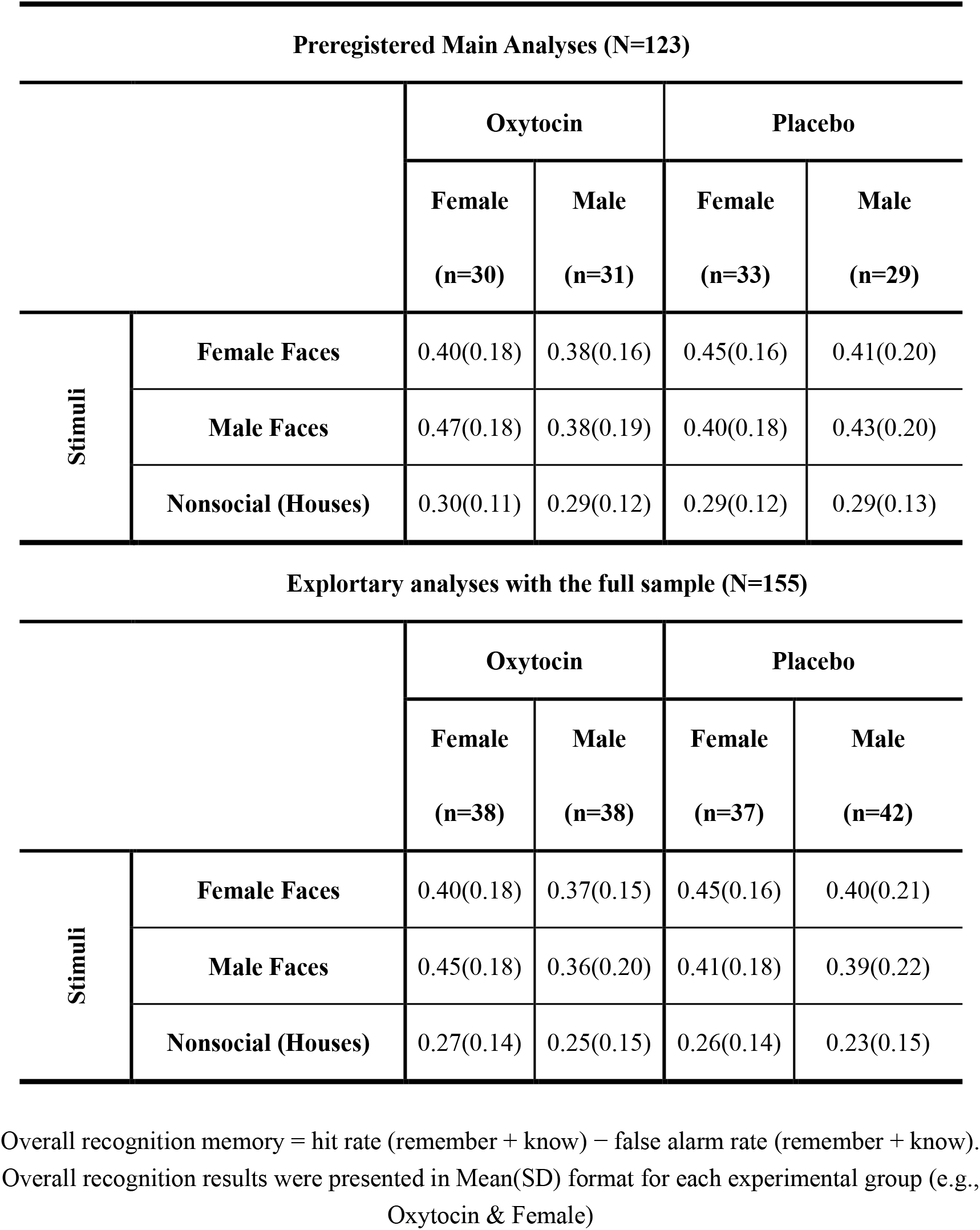
Memory Performance (i.e., Overall Recognition) of Post-encoding Study1.

We firstly analyzed data excluding particpants with low retrieval performance (123 out of 155 participants were analyed). Our analysis revealed a significant three-way interaction among the gender of the faces, the gender of the participants, and oxytocin administration (F_(1,119)_=4.86, *p*=0.029, partial η2 =.039; **Figure 2A**). Specifically, among female participants, a significant interaction was observed between the gender of the faces and oxytocin administration (F_(1,61)_=7.31, *p*=0.009, partial η2 = .107). Female participants who received oxytocin after encoding exhibited enhanced memory (t_29_=2.25, *p*=0.03, Cohen’s d=0.41) for male faces (M_female_=0.470, SD_male_=0.177) as opposed to female faces (M_male_=0.402, SD_male_=0.184). Conversely, female participants who received the placebo did not show this memory advantages for opposite-gender faces (M_female_=0.453, SD_female_=0.158; M_male_=0.398, SD_male_=0.178; t_32_=1.63, *p*=0.11, Cohen’s d=0.285). No similar gender of the faces and oxytocin administration interaction was found among male participants (F_(1,58)_=0.26, *p*=0.61, partial η2 =0.004).

**Figure 2.**
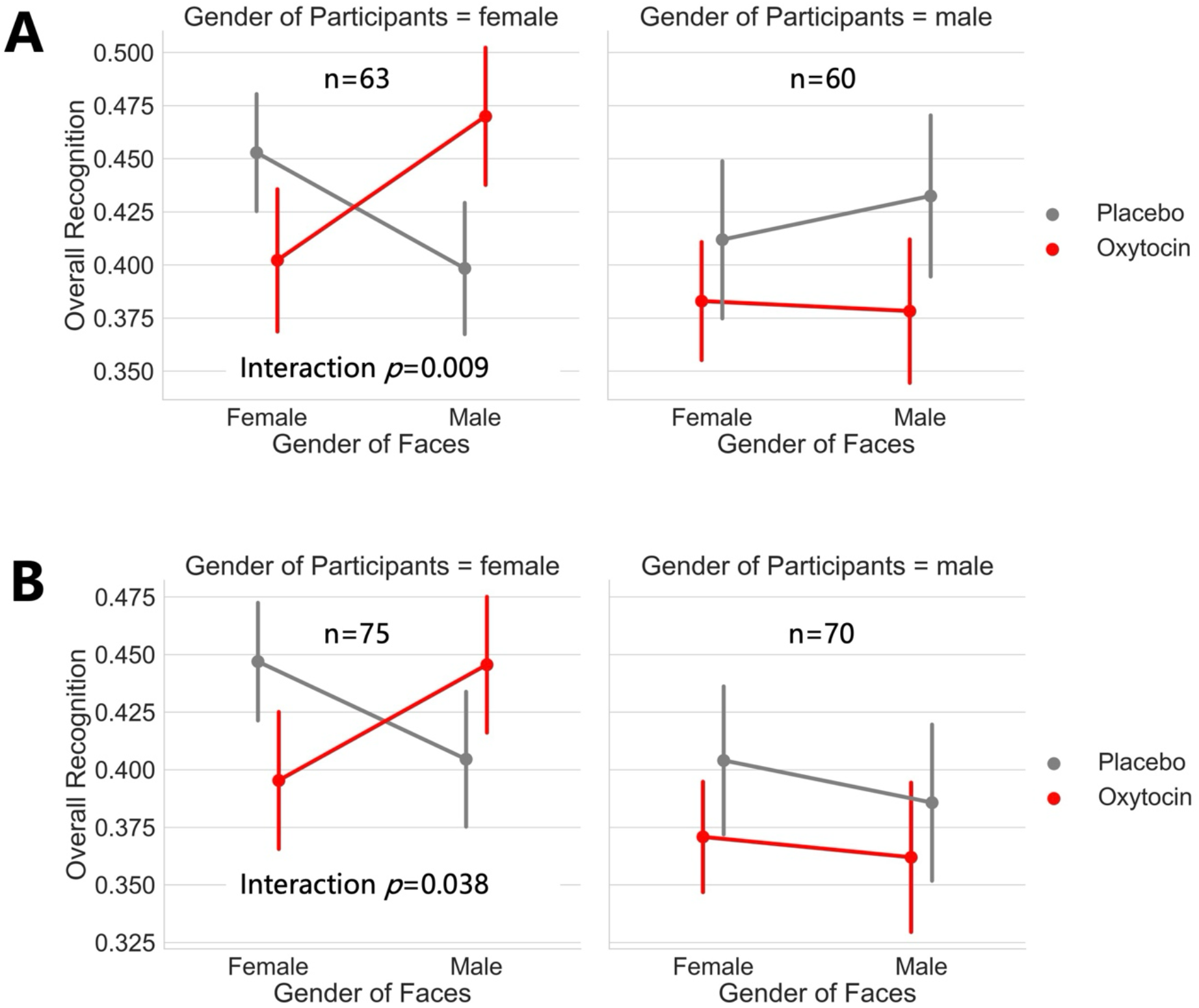
Oxytocin after encoding enhances recognition of male faces among female participants. **(A) Preregistered main analyses.** Examining overall recognition performance in a sample of 123 participants, we found a significant three-way interaction between face gender, participant gender, and oxytocin administration (F_(1,119)_=4.86, *p*=0.029, partial η2 =.039). Specifically, in female participants, a significant interaction was observed between the face gender and oxytocin administration (F_(1,61)_=7.31, *p*=0.009, partial η2 = .107). Females receiving oxytocin post-encoding demonstrated significantly improved memory for male faces over female faces (t_29_=2.25, *p*=0.03, Cohen’s d=0.41). **(B) Exploratory Analyses:** Using the full sample (N = 155), we replicated the interaction between the gender of the faces and oxytocin administration among female pariticpants (F_(1,73)_=4.46, *p*=0.038, partial η2 = .058) from the preregistered analyses.

Our exploratory analyses, which included all 155 participants regardless of their retrieval performance. Although we did not replicated the significant three-way interaction among the gender of the faces, the gender of the participants, and oxytocin administration (F_(1,151)_=1.71, *p*=0.194, partial η2 =.011; **Figure2B**), among female participants, we replicated a significant interaction with identifcal direction was observed between the gender of the faces and oxytocin administration (F_(1,73)_=4.46, *p*=0.038, partial η2 = .058). This exploratory analysis suggests that the reported memory enhancement effects did not depend on our pre-defined specific participant exclusion procedures.

### Social placebo effects or different approachability during encoding cannot explain memory enhancements

Subsequent analyses were conducted to rule out alternative explainations for the observed enhancement effect of oxytocin on memory consolidation of human faces. Firstly, we considered the possibility that participants could discern whether they had received oxytocin or a placebo, potentially leading to biased rehearsal of specific stimulus types between the day1 and day2 due to the social placebo effect^23^, and ultimately enhancing social memories. To assess this, participants were asked to identify the treatment (oxytocin or placebo) they believed they had received during day1 at the end of the entire experimental procedure. Of the participants, 28 out of 61 (approximately 46%) in the oxytocin group and 26 out of 62 (approximately 42%) in the placebo group presumed they were in the oxytocin group (χ² = 0.196, *p* = 0.65), indicating a general confusion about their actual experimental conditions. Among female participants who received oxytocin, whether they believed they had received oxytocin or not did not affect their memories for male faces (t_28_=-0.157, *p*=0.876, Cohen’s d=-0.058) or female faces (t_28_=-0.721, *p*=0.477, Cohen’s d=-0.266). As exploratory analyses for the social placebo effect, we explored the interaction between gender of face and perceived oxytocin administration. There was no significant interaction (F_(1,61)_=2.725, *p*=0.104, partial η2 = 0.043), suggesting that the social placebo effect can not explain the memory enhancement effect.

Secondly, we explored whether inherent group differences (e.g., different levels of efforts) during encoding could account for the differential memory performances observed on the day2. This was investigated by analyzing approachability ratings of faces during the encoding phase on a trial-by-trial basis. Due to software and computer errors, the approachability ratings for 7 out of a total of 123 participants were not recorded properly and were therefore excluded from the analyses. Unlike memory performance, we did not find a significant three-way interaction among the gender of the faces, the gender of the participants, and oxytocin administration on approachability (F_(1,112)_=2.87, *p*=0.093, partial η2 =.025). When we restricted our analyses to female participants, we found that participants in both the oxytocin (*p*=0.002) and placebo groups (*p*<0.001) reported a higher likelihood of approaching faces of the same gender (i.e., female participants were more likely to approach female faces). This was expected to lead to equal memory advantages for female faces in both groups. However, our experimental evidence showed that post-encoding administration of oxytocin reversed this pattern, resulting in stronger memory for opposite-gender faces (i.e., male faces) during the 24-hour delayed memory retrieval.

### No stimuli-specific or valence-specific memory enhancement due to oxytocin

As pre-registered, we preformed planned analyses to examine whether oxytocin could (1) selectively enhance memories of specific types of stimuli (i.e., face) or (2) specific valences (i.e., enhance positive and suppress negative memories). We restricted our analyses to 123 participants who passed the data excludion procedure. Results showed that: (1) although faces were better rememberd than house (M_face_=0.41, SD_face_=1.54; M_house_=0.29, SD_house_=0.11; t_122_=8.234, *p*<0.001, Cohen’s d=0.742), this enhancement did not vary with oxytocin administration (F_(1,121)_=0.628, *p*=0.43, partial η2 =0.005). (2) Although the valence of faces modulated retrieval performance (M_postive_=0.420, SD_postive_=0.194; M_neutral_=0.308, SD_neutral_=0.207; M_negative_=0.517, SD_negative_=0.211; F_(2,244)_ = 49.56, *p* < 0.001, partial η2 =0.289), this difference did not vary with oxytocin administration (F_(2,242)_ = 0.02, *p* = 0.976, partial η2 <0.001). This indicates that oxytocin uniformly affected face memories across different valence categories, suggesting that the enhanced memory performance cannot be attributed to selective improvement of certain valence categories.

### Enhanced memories cannot be explained by the oxytocin-affected memory retrieval

To further rule out the possibility that enhanced memory performance reported in study1 is the result of long-lasting effects (i.e., 24 hours) of oxytocin on retrieval, we administrated oxytocin 30 minutes before retrieval (23.5h after the intial study session) in Study2 to directly examine its effects on memory retrieval. Study 2 was a preregistered, randomized, double-blind, placebo-controlled trial involving heterosexual participants. Despite differences in the timing of oxytocin administration, both studies were identical in experimental procedures, stimuli, trial structure, memory assessment methods, and utilized the same set of computers and experimenters. A total of 139 patitipants (70 female and 69 male) were tested, with 121 passing the data exclusion criteria (**Table 2**). Firstly, we attempted to replicate the three-way interaction among the gender of the faces, the gender of the participants, and oxytocin administration in Study1 but failed to find the significant three-way interaction (F_(1,117)_=0.69, *p*=0.40, partial η2 =0.006). Even among female participants, there was no significant interaction between the gender of the faces and oxytocin administration (F_(1,59)_=0.04, *p*=0.839, partial η2 =<0.001). Next, we preformed planned analyses to exaime whether oxytocin before retrieval could (1) selectively enhance face memories or (2) modulate face memories in a valence-specific way. Results showed that: (1) there was no internaction between type of stimuli and oxytocin administration (F_(1,119)_=1.246, *p*=0.266, partial η2 =0.01). (2) Although the valence of faces modulated retrieval performance (M_postive_=0.417, SD_postive_=0.208; M_neutral_=0.320, SD_neutral_=0.202; M_negative_=0.546, SD_negative_=0.210; F_(2,240)_ = 60.73, *p* < 0.001, partial η2 =0.336), this difference did not vary with oxytocin administration (F_(2,238)_ = 0.511, *p* = 0.60, partial η2 =0.001).

**Table 2.** Memory Performance (i.e., Overall Recognition) of Before Retrieval Study2.

| Preregistered Main Analyses (N=121) |  |  |  |  |  |
| --- | --- | --- | --- | --- | --- |
|  |  | Oxytocin |  | Placebo |  |
|  |  | Female | Male | Female | Male |
|  |  | (n=30) | (n=30) | (n=31) | (n=30) |
| Stimuli | Female Faces | 0.45(0.18) | 0.40(0.20) | 0.47(0.15) | 0.42(0.20) |
|  | Male Faces | 0.45(0.20) | 0.39(0.22) | 0.49(0.18) | 0.36(0.20) |
|  | Nonsocial Houses | 0.45(0.16) | 0.27(0.12) | 0.48(0.14) | 0.30(0.16) |

Overall recognition memory = hit rate (remember + know) – false alarm rate (remember + know). Overall recognition results were presented in Mean(SD) format for each experimental group (e.g., Oxytocin & Female)

Although oxytocin had no impact on memory retrieval, we also analyzed the social placebos effect. Of the Study2 participants, 21 out of 60 (approximately 35%) in the oxytocin group and 17 out of 61 (approximately 28%) in the placebo group presumed they were in the oxytocin group (χ² = 0.714, *p* = 0.398), indicating a general confusion about their actual experimental conditions. Among female participants who received oxytocin, whether they believed they had received oxytocin or not did not affect their memories for male faces (t_28_=1.595, *p*=0.122, Cohen’s d=0.658) or female faces (t_28_=1.552, *p*=0.132, Cohen’s d=0.641). As exploratory analyses for the social placebo effect, we explored the interaction between gender of face and perceived oxytocin administration. There was no significant interaction (F_(1,59)_=1.218, *p*=0.274, partial η2 = 0.02). In summary, we did not find evidence that before-retrieval oxytocin adminstration could modulate memory performance in any format in Study 2, suggesting that the reported significant memory enhancement effect in Study 1 cannot be explained by long-lasting effects of oxytocin on retrieval.

## Discussion

This study demonstrates that administering oxytocin immediately after encoding significantly enhances memory consolidation of human faces, with effects varying by the gender of both the participants and the faces remembered. Specifically, oxytocin administration resulted in a roughly 20% increase in recognition of male faces by female participants (i.e., from 0.40 to 0.47). Previous studies have reported a robust own-gender bias in face memory, where female participants typically recall more female than male faces^24^. Our findings suggest that oxytocin can reverse this bias during the consolidation phase, supporting theories that link oxytocin to social bonding effects^2,25^.

Given that oxytocin was administered post-encoding, its primary influence likely pertains to memory consolidation rather than altered encoding or retrieval processes. This conclusion is further supported by additional analysis of self-reported approachability during encoding and by Study 2, which directly assessed oxytocin’s impact on retrieval. While the mechanisms by which oxytocin enhances social memory consolidation are not fully understood, our results indicate that its effects are mechanistically distinct from those induced by exercise^12^, caffeine^13^, emotion^15^, and arousal^16^. Unlike typical memory modulation during consolidation mentioned above, which is dependent on the participant’s state and non-selective towards specific memories, our evidence shows that oxytocin-induced memory consolidation can be selective and dependent on participant traits such as gender.

The most plausible explanation for the selective rather than overall enhancement of memory is that oxytocin facilitates social memory consolidation by promoting the spontaneous reactivation of faces during awaking rest^26,27^ or sleep^28^. It is still unclear whether the effects of oxytocin require sleep or prolonged consolidation periods, or if they might appear during or shortly after administration—for instance, within 6 hours—since our study only assessed memory 24 hours post-encoding. Moreover, the specific brain networks involved in oxytocin-induced memory consolidation enhancement for human faces remain indeterminate from our data. Future research should explore this by recording spontaneous post-encoding neural activity^14,29^ and employing computational decoding techniques to analyze this activity^30^. At the same time, both peripheral and central oxytocin levels during memory consolidation could be monitored and correlated with neural activity^31^. Such monitoring could provide insights into the temporal dynamics and mechanistic underpinnings of oxytocin’s influence on memory consolidation.

We tested three pre-registered hypotheises and found no evidence to support the stimuli and valence-specific enhancement of memory consolidation. Initially, Rimmele et al. demonstrated that oxytocin, administered before memory encoding, selectively improves recognition memory for faces (but not for non-social stimuli) when tested 24 hours later^9^. It is crucial to note that since the oxytocin was administered prior to learning and memory was assessed twenty-four hours later, it may have simultaneously enhanced memory encoding and consolidation of face memories, resulting in improved long-term memory retention. In our study, we employed both post-encoding and pre-retrieval design (i.e., oxytocin administered after memory encoding in Study1 and before retrieval in Study2) and an episodic memory task closely mirroring Rimmele’s study. Nevertheless, we observed no similar selective memory enhancements in either male or female participants (note that Rimmele’s study did not include female participants). We propose that the effects reported in Rimmele’s study are more likely attributable to enhanced memory encoding of faces due to oxytocin, rather than consolidation. Secondly, our findings revealed no evidence that oxytocin-induced enhancement of social memory depends on the valence. Aligned with the social adaptation model of oxytocin^2^ and corresponding neuroimaging findings in humans^32^, our original hypothesis posited that oxytocin could enhance the consolidation of positive social memories while reducing that of negative ones. Yet, this hypothesis was not corroborated by empirical evidence, suggesting that oxytocin’s effects might differ across memory encoding and consolidation phases. Enhanced encoding appears more closely related to the sensitivity of the information, whereas enhanced consolidation may be associated with the neural reactivation process.

In conclusion, our results provide solid evidence from two preregistered, randomized, double-blind, placebo-controlled trials with a large sample of heterosexual participants (N=294) that oxytocin can affect social memory consolidations in humans. Overall, our findings extend the understanding of oxytocin’s effect on social memories of in humans, broaden the knowledge of the so-called love hormone’s, oxytocin’s, role in social cognition^2,3,25^, and highlight its potential as a novel pharmacological strategy for selectively enhancing memory consolidation in humans. Our experiment thus serves as a proof-of-principle study that could inspire future applications of oxytocin to reverse social memory deficits in clinical populations such as those with autism^4^ and schizophrenia^33^.

## Methods

### Resource Availability

#### Materials Availability

The scripts and code used in this study for for data collection are accessible at https://osf.io/5emcf/?view_only=b365566fe0cc4ec0b7798f197d84a2ee.

#### Data and Code Availability

The datasets and code used in this study for data analysis are available at https://osf.io/5emcf/?view_only=b365566fe0cc4ec0b7798f197d84a2ee.

#### Preregistration Links

Study1 was preregistered as the project titled: *The Effects of Oxytocin on Episodic Memory Consolidation in Humans* (Link: https://doi.org/10.17605/OSF.IO/8PXMR).

#### Study2 was preregistered as the project titled: *The Effects of Oxytocin on Episodic Memory Retrieval in Humans* (Link: https://doi.org/10.17605/OSF.IO/MU3BF)

#### Lead Contact

Further information and requests for resources should be directed to and will be fulfilled by the Lead Contact, Wei Liu.

### Participants

We examined 294 nonsmoking, healthy, heterosexual individuals without any psychiatric, neurological, or medical illnesses. From the initial sample, we excluded data from 50 participants whose recollection, familiarity, or overall recognition scores (any of the three memory measures) were below zero, indicating they had failed to encode the material properly during the memory encoding phase. Consequently, 244 participants were included in the preregistered main analyses: ages ranged from 18 to 28 years (mean: 20.30, SD: 1.74), comprising 120 males (ages 18-28, mean: 20.38, SD: 1.93) and 124 females (ages 18-26, mean: 20.23, SD: 1.55). The oxytocin group included 61 males and 60 females (total n=121, ages 18-26, mean: 20.31, SD: 1.69), while the placebo group included 59 males and 64 females (total n=123, ages 18-28, mean: 20.29, SD: 1.80). The study was approved by the Institutional Review Committee of Central China Normal University. All subjects provided written informed consent and received partial compensation. Participants on medication or reporting abnormal sleep-wake cycles, known allergies, acute or chronic nasal disorders, or nasal congestion were excluded. Participants were instructed to avoid caffeinated or alcoholic beverages during the experiment and to maintain a regular sleep-wake cycle.

#### Study 1

A total of 123 nonsmoking, healthy, heterosexual participants received the administration immediately after completing the encoding phase on the first day of the experiment. Participants’ ages ranged from 18 to 28 years (mean: 20.54, SD: 1.85), including 60 males (ages 18-28, mean: 20.60, SD: 2.51) and 63 females (ages 18-24, mean: 20.48, SD: 1.65). The oxytocin group comprised 31 males and 30 females (total n=61, ages 18-26, mean: 20.43, SD: 1.86), and the placebo group comprised 29 males and 33 females (total n=62, ages 18-28, mean: 20.63, SD: 1.85). Data from 32 participants were excluded from the analysis.

#### Study 2

A total of 121 nonsmoking, healthy, heterosexual participants received the administration immediately after completing the encoding phase on the first day of the experiment.

Participants’ ages ranged from 18 to 26 years (mean: 20.07, SD: 1.60), including 60 males (ages 18-25, mean: 20.15, SD: 1.78) and 61 females (ages 18-26, mean: 19.98, SD: 1.41). The oxytocin group comprised 30 males and 30 females (total n=60, ages 18-24, mean: 20.18, SD: 1.50), and the placebo group comprised 30 males and 31 females (total n=61, ages 18-26, mean: 19.95, SD: 1.70). Data from 18 participants were excluded from the analysis.

### Experimental Procedures

On Day 1, participants incidentally encoded images of social (i.e., human faces) and non-social (i.e., houses) stimuli (see *Memory Task-Encoding* below). The timing of administration was immediate—post-encoding in Study 1 and pre-retrieval in Study 2. After 24 hours (i.e., Day 2), we unexpectedly evaluated participants’ memory retention using the remember/know (RK) paradigm (see *Memory Task-Retrieval* below). Study 1 and Study 2 were identical except for the timing of oxytocin administration.

### Double-Blind and Randomization

The study was conducted as a randomized, double-blind, placebo-controlled intervention. We used block randomization, where each participant was randomly assigned to one of two equally sized, predetermined blocks (oxytocin or placebo). The random number list used to create these blocks was generated using web-based applications. To maintain the double-blind protocol, we used labels A and B to represent oxytocin and placebo during data collection. The relationship between A/B and oxytocin/placebo was not known to the on-site experimenters until the final data analysis phase. Additionally, participants were asked to identify the treatment (oxytocin or placebo) they believed they had received at the end of the experimental procedure to statistically assess the success of the double-blind operation.

### Administration of Oxytocin

Recent studies have shown that intranasal administration of neuropeptides, such as vasopressin, enables peptides to enter the central nervous compartment directly, providing a useful method to study the effects of the human neuropeptide oxytocin on the central nervous system. No adverse side effects were reported in our experiments. Subjects received a single dose of 24 IU of oxytocin (Syntocinon Spray; Novartis; three puffs per nostril, each puff containing 4 IU of oxytocin) intranasally or a placebo (containing all inactive ingredients except the neuropeptide). Oxytocin was administered immediately after encoding in Study 1 (i.e., post-encoding condition) and 30 minutes before retrieval in Study 2 (i.e., before-retrieval condition).

### Memory Task

#### Stimuli

The experiment used a total of 90 social stimuli (i.e., human faces) and 90 non-social stimuli (i.e., houses). Using materials from the Chinese Academy of Sciences’ Sinicized Face Emotion Image System, we randomly selected 90 gray-scale facial stimuli (45 men, 45 women) from 600 images, including 30 faces each with positive, neutral, and negative emotions. Of these, 60 faces (30 men, 30 women) were used as encoding images, comprising 20 images each of positive, neutral, and negative emotions. The remaining 30 faces (15 men, 15 women), comprising 10 images each of positive, neutral, and negative emotions, were used as interference images. For non-social stimuli, we selected 90 images (30 houses, 30 castles, 30 offices) from pixabay.com to assess non-social memory. These stimuli were presented in an oval mask on a black background. Like the social stimuli, 60 images (20 houses, 20 castles, 20 offices) were used for encoding, and 30 images (10 houses, 10 castles, 10 offices) were used as interference images.

#### Encoding

On the first day, participants engaged in an approachability rating task. They were presented with 60 faces and 60 non-social stimuli in two modules, each containing 60 stimulus images, with a 20-second interval between the two modules. Each trial began with the presentation of a picture for 3500 ms, followed by a self-paced rating on an approachability scale ranging from 1 (not at all) to 5 (very approachable). The approachability rating served as a cover story for the memory encoding task, ensuring proper encoding by requiring participants to indicate how much they would like to approach the presented stimulus. The encoding time was constant for all stimuli (3500 ms), as participants rated approachability after the stimulus disappeared from the monitor. An interstimulus interval (ISI) of 500 ms separated each trial.

#### Retrieval

We unexpectedly evaluated participants’ memory retention using the remember/know (RK) paradigm. On the second day (i.e., 24 hours later), the faces and non-social stimuli encoded the day before were subjected to unanticipated recognition memory tests. The 60 face pictures and 60 non-social stimulus pictures from the previous day were presented again, along with 30 new face pictures and 30 new non-social stimulus pictures, randomly mixed in two blocks, for a total of 180 stimulus pictures. Participants had to indicate whether they remembered (accurately recalling several details of the image), knew (recognized the image but could not recall details), or found the image new (not seen at the time of encoding) by pressing one of three response keys. Each trial began with the presentation of a picture for 3500 ms, followed by a self-paced rating on the RK paradigm (remember, know, new). An ISI of 500 ms separated each trial. Before the formal memory retrieval phase, participants practiced the responses using a new set of stimuli (including cartoon faces and buildings unrelated to the main experiments). The practice phase, including encoding and retrieval, lasted for about 3-5 minutes to ensure participants clearly understood the difference between “remember” and “know.”

### Statistical Analysis

#### Memory performance analyses

Overall recognition memory accuracy was assessed by subtracting the overall false alarm rate from the overall hit rate. Recollection and familiarity were estimated separately using remember and know responses. Recollection was measured by subtracting the proportion of new items receiving a remember response (false alarms with a remember response) from the proportion of old items receiving a remember response (hits with a remember response). Familiarity was calculated as the probability of responding "know" to an item, given that the item did not receive a remember response, corrected for false alarms [i.e., (hit rate know / (1 - hit rate remember)) - (false alarm rate know / (1 - false alarm rate remember))]. In our main analyses, overall recognition was used as the primary memory measure to investigate the effects of oxytocin on long-term memory: Overall recognition memory = hit rate (remember + know) − false alarm rate (remember + know). Recollection = hit rate (remember) − false alarm rate (remember). Familiarity = (hit rate know/(1 − hit rate remember)) − (false alarm rate know/(1 − false alarm rate remember)). The statistical analysis relied on ANOVA, including a three-factor mixed ANOVA with two intergroup factors, treatment (oxytocin vs. placebo) and the gender of the participants (male vs. female), and a repeated measure factor, the gender of the faces (male vs. female). Additionally, a two-factor mixed analysis of variance, with an intergroup factor treatment (oxytocin vs. placebo) and a repeated measure factor, the gender of the faces (male vs. female), was used to investigate specific changes in certain genders. Significant analyses of variance, main effects, or interactions were identified using two-way and three-way ANOVAs, paired sample t-tests, and independent sample t-tests. All tests were bilateral, and the significance level was set at p = 0.05. All statistical analyses were performed using JASP (https://jasp-stats.org/)

#### Social placebo effect and approachability rating analyses

Aside from the placebo control at the experimental procedure level, such as the randomized, double-blind, placebo-controlled design, we performed additional analyses to further rule out placebo effects. First, we used a chi-square test to analyze participants’ guesses about whether they were in the treatment group (i.e., oxytocin group). Second, we analyzed memory performance based on participants’ guessed treatment instead of the actual treatment to see whether believing they belonged to the oxytocin group could also produce significant experimental effects.

We analyzed approachability ratings during encoding to rule out the possibility that different encoding processes could explain the memory enhancement effect. We performed a three-factor mixed ANOVA with two intergroup factors, treatment (oxytocin vs. placebo) and the gender of the participants (male vs. female), and a repeated measure factor, the gender of the faces (male vs. female), and a two-factor mixed analysis of variance with an intergroup factor treatment (oxytocin vs. placebo) and a repeated measure factor, the gender of the faces (male vs. female) on approachability ratings. If the changes in approachability mirrored the memory performance, it would suggest that different encoding phases contribute to the final memory performance.

## Author contribution statement following the CRediT Taxonomy

**W.L**: Conceptualization: Lead; Funding acquisition: Lead; Project administration: Lead; Supervision: Lead; Validation: Lead; Visualization: Lead; Writing – original draft: Lead; Writing – review & editing: Lead. **J.S.L**: Conceptualization: Supporting; Data Curation:Lead; Software:Lead; Investigation: Lead; Formal Analysis: Lead; Writing – original draft: Supporting; Writing – review & editing: Supporting. **Z.Y.C**: Data Curation: Supporting; Software: Supporting; Investigation: Supporting; Formal Analysis: Supporting; Writing – original draft: Supporting

## Acknowledgement

W.L was supported by the National Natural Science Foundation of China (grant No. 32300879), Humanities and Social Sciences Fund, Ministry of Education (grant No. 22YJCZH109), Natural Science Foundation of Hubei (grant No. 2022CFB793), Open Research Fund of the State Key Laboratory of Cognitive Neuroscience and Learning (CNLYB2103), Open Research Fund of the Key Laboratory of Adolescent Cyber Psychology and Behavior (grant No. CCNUCYPSYLAB2022B10), and the Major Program of the National Social Science Foundation of China (grant No. 22&ZD187).

